# Genetic, environmental and intrinsic determinants of the human antibody epitope repertoire

**DOI:** 10.1101/2021.12.07.471553

**Authors:** Sergio Andreu-Sánchez, Arno R. Bourgonje, Thomas Vogl, Alexander Kurilshikov, Sigal Leviatan, Angel J. Ruiz Moreno, Shixian Hu, Trishla Sinha, Arnau Vich Vila, Shelley Klompus, Iris N. Kalka, Karina de Leeuw, Suzanne Arends, Iris Jonkers, Sebo Withoff, Lifelines cohort study, Elisabeth Brouwer, Adina Weinberger, Cisca Wijmenga, Eran Segal, Rinse K. Weersma, Jingyuan Fu, Alexandra Zhernakova

## Abstract

Phage-displayed immunoprecipitation sequencing (PhIP-Seq) has successfully enabled high-throughput profiling of human antibody profiles. However, a comprehensive overview of environmental and genetic determinants shaping human adaptive immunity is currently lacking. In this study, we aimed to investigate the effects of genetic, environmental and intrinsic factors on the variation in human antibody repertoires. We characterized serological antibody repertoires against 344,000 peptides using PhIP-Seq libraries from a wide range of microbial and environmental antigens in 1,443 participants from a population cohort. We demonstrate individual-specificity, temporal consistency and co-housing similarities in antibody repertoire. Genetic analyses showed involvement of the HLA, IGHV and FUT2 regions. Furthermore, we uncovered associations between 48 phenotypic factors and 544 antibody-bound peptides, including age, cell counts, sex, smoking behavior and allergies, among others. Overall, our results indicate that human antibody epitope repertoires are shaped by both host genetics and environmental exposures and highlight unique signatures of distinct phenotypes and genotypes.

## Introduction

The adaptive immune system encompasses an extremely complex group of biological processes that orchestrate responses to invading pathogens in all jawed vertebrates (*gnathostomes*), including humans (Cooper and Alder, 2006). The adaptive immune system’s capacity to recognize, adapt to and remember a wide variety of threats is determined by highly polymorphic genetic structures that encode receptors able to interact with complex structures known as antigens that most commonly represent amino acid sequences (epitopes) from foreign proteins (Cooper and Alder, 2006). Antibodies are the key effector molecules in the human adaptive immune system and are responsible for humoral immunity. Each individual’s antibody epitope repertoire is characterized by a high degree of versatility and adaptability and is continuously altered during their lifetime, with host genetics and environmental factors being the main contributors. Antibody repertoires determine the fate of the immune response against pathogens and the development of autoimmunity or allergies, and they have garnered special attention because they can be used to study herd immunity acquisition (Burkholder et al., 2017). In an adult human, there are around 10^10^–10^11^ B-lymphocytes, each expressing a unique B-cell receptor (BCR) (a non-soluble antibody form) that identifies a molecular pattern (Ganusov and De Boer, 2007). The antigenic diversity of BCRs is the net result of the high diversity and somatic rearrangements of V(D)J gene segments, insertion and deletion (indel) of nucleotides and subsequent somatic hypermutation (SHM) to increase antigen affinity and specificity (Hoehn et al., 2016).

To gain more insight into antibody–antigen interaction, efforts have been made to directly sequence the BCR (Galson et al., 2020; Goldstein et al., 2019) and to directly infer it from single-cell transcriptomic sequencing (Lindeman et al., 2018). Although this methodology provides information on the potential for generation of immune responses against yet unknown antigens, it does not directly link BCR sequences to the exact nature of antigenic epitopes. In addition, in terms of scaling, it is limited to just a small proportion of the immense number of these receptors (Kim and Park, 2019). On the other hand, antibody-binding analysis, such as peptide microarrays (Atak et al., 2016; Yu et al., 2017) or enzyme-linked immunosorbent assay (ELISAs), enable the determination of antibody seroprevalence against selected antigens. While easily implemented for a limited set of antigens, these methodologies have been difficult to scale up to thousands of antigens in a large population. Phage-displayed immunoprecipitation sequencing (PhIP-Seq) is an immuno-precipitation–based sequencing technique that enables quantification of antigen peptides that are displayed as phage libraries, which subsequently react with human antibodies, and antibody-bound phages are eventually sequenced to obtain an ‘immunological fingerprint’ of an individual’s antibody repertoire. PhIP-Seq has been described previously (Larman et al., 2011; Mohan et al., 2018) and has been successfully applied to characterize autoimmune antibody prevalence in patients with multiple sclerosis, type 1 diabetes and rheumatoid arthritis (Larman et al., 2013; Román-Meléndez et al., 2021), the human virome (Eshleman et al., 2019; Finton et al., 2014; Mina et al., 2019; Shrock et al., 2020; Xu et al., 2015), the widespread presence of antibodies against virulence factors (Angkeow et al., 2021; Vogl et al., 2021) and the gut microbiome (Vogl et al., 2021). However, no comprehensive study has been carried out to date that identifies the environmental, intrinsic, lifestyle and genetic factors that determine antibody generation against antigen exposures in the general population.

In this work, we set out to uncover the antibody epitope repertoire in a deeply phenotyped population cohort from the northern part of the Netherlands, Lifelines-DEEP (LLD) (Tigchelaar et al., 2015). We used the PhIP-Seq libraries described in (Vogl et al., 2021) and (Leviatan et al., in prep) to characterize 344,000 peptide antigens related to: (1) microbes (including human gut microbiota, probiotic strains, pathobionts, antibody-coated species and virulence factors from the virulence factors database (VFDB)), (2) the immune epitope database (IEDB) (Vita et al., 2015), (3) proteins from allergen databases and (4) bacteriphages.

Leveraging the rich metadata available for this deeply phenotyped cohort (including imputed genotypes, gut microbiota shotgun sequencing, clinical blood tests (immune, metabolic and autoimmune markers), family information, lifestyle and self-reported diseases and allergy questionnaires) alongside the PhIP-Seq data allowed us to establish key genetic and environmental factors shaping the human antibody epitope repertoires.

## Results

### Antibody-bound peptide repertoires are personalized, linked to shared environments (co-housing) and time-dependent

We interrogated a total of 344,000 peptides in 1,778 samples from 1,437 individuals (for 341 of whom we have data at two time points 4-years apart) from a northern Dutch population cohort (Tigchelaar et al., 2015) [**Fig 1A**].

**Figure 1.**
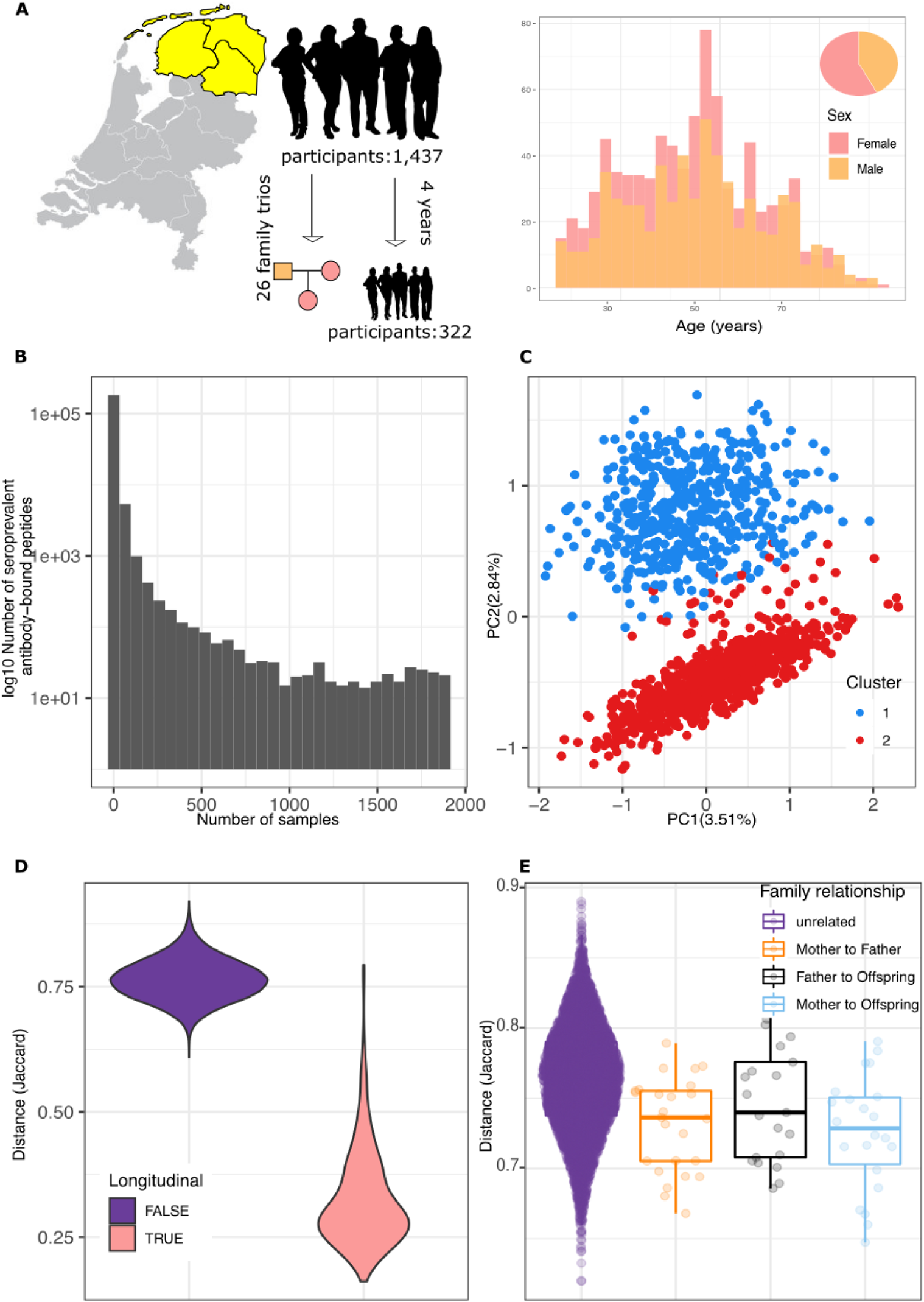
PhIP-Seq antibody-bound peptide profiles of 1,443 individuals representative of the Dutch population. **A.** Cohort characteristics. Lifelines-Deep is a population cohort from Northern Netherlands. In this work, we performed PhIP-Seq in 1,443 participants (including 26 trio families), 322 of whom have data from a second time point after 4 years. Other data layers include phenotypes (questionnaires and clinical measurements), genetics (imputed microarrays) and microbiome (bacterial taxonomic quantification). There is a higher proportion of females within the participants (57%). The age distribution is slightly left skewed, with a mean of 44.5 years (no significant sex differences). **B.** Prevalence of antibody-bound peptides in the population. X-axis depicts seroprevalence. Y-axis is the number of antibody-bound peptides with a given seroprevalence. **C.** Principal component analysis identified two clusters (color represents cluster labels after 2-means clustering) related to PC2, which is mainly driven by CMV peptides. **D.** Longitudinal samples taken 4 years apart in 322 participants are more closely related to the individual’s baseline than to other participants. **E.** 26 family trios show a lower distance between their antibody-bound peptide profiles than unrelated participants.

After immunoprecipitation with protein A/G (binding primarily IgG antibodies (Vogl et al., 2021)) and sequencing, we detected an enrichment of sequenced reads (compared to a null distribution without immunoprecipitation) of 175,242 (antibody-bound) peptides in at least one participant (average number of peptides bound per person = 1,168, range = 3–3,161) (see Methods). Peptide seropositivity was defined as a presence/absence binary score (enriched/not enriched) that was used for all subsequent analyses. Most antibody-bound peptides showed low seroprevalence, indicating the individual-specificity of the antibody epitope repertoire [**Fig 1B**]. Based on peptide sequence identity and prevalence (see Methods for details), we chose 2,815 peptides for further analyses [**Supplementary Table 1.1**].

The large variability in the antibody-bound peptide enrichment profile can be seen through a principal component analysis (PCA), where the amount of variability recovered by the first 10 components was just 15.5% and 709 components were needed to retrieve 90% of the total antibody-bound peptide variability [**Fig 1C**]. Despite the relatively low variability accounted for by the first two PCs (6.3%), we observed two clusters in PC2 that were driven by cytomegalovirus (CMV)-related antibody-bound peptides (K-means, k = 2) [**Figure 1A**] (removal of these peptides resolves PC2 clustering, **Supplementary Fig 1A**).This is consistent with a previous observation that nearly 50% of the Dutch adult population are seropositive for this herpesvirus (Korndewal et al., 2015). In contrast, PC1 was highly related to the number of seropositive peptides (affine linear model R^2^ = 72%). In a permutational multivariate analysis of variance (PERMANOVA), the person-to-person antibody-bound peptide repertoire dissimilarity showed effects of age (R^2^=1.4%), lifestyle phenotypes (smoking R^2^=0.18%), blood measurements (cholesterol R^2^=0.12%) and blood cell counts (lymphocytes relative abundance, R^2^=0.16%), among many others (while correcting for age, sex and sequencing plate) [**Supplementary Table 2.1**].

In agreement with previous reports, we observed temporal consistency in the antibody-bound peptide repertoire (Angkeow et al., 2021; Vogl et al., 2021) for the 322 LLD participants who were followed-up after 4 years. We observed that the distance between samples taken from the same individuals 4-years apart were on average lower than the distance of unrelated individuals (p < 5×10^−4^; 2,000 label permutations) [**Figure 1D**]. Overall, the distance between baseline and follow-up was not associated with baseline age or sex. The temporal consistency of antibody-bound peptides showed a binomial distribution, with most peptides consistent between timepoints while a subset others tended to change [**Supplementary Fig 2B**][**Supplementary Table 1.1**]. This change was more often a loss of enrichment rather than gain, and this difference could not be directly attributed to a batch effect (Wilcoxon test, p = 0.45). This highlights that the time elapsed since antigen encounter might be a determining factor for the detection of antibody-bound peptide enrichment, which is in agreement with humoral studies showing that the prevalence of antibodies seemed to fade over time (Erles et al., 1999; Hendrikx et al., 2011; Kontio et al., 2012).

**Figure 2.**
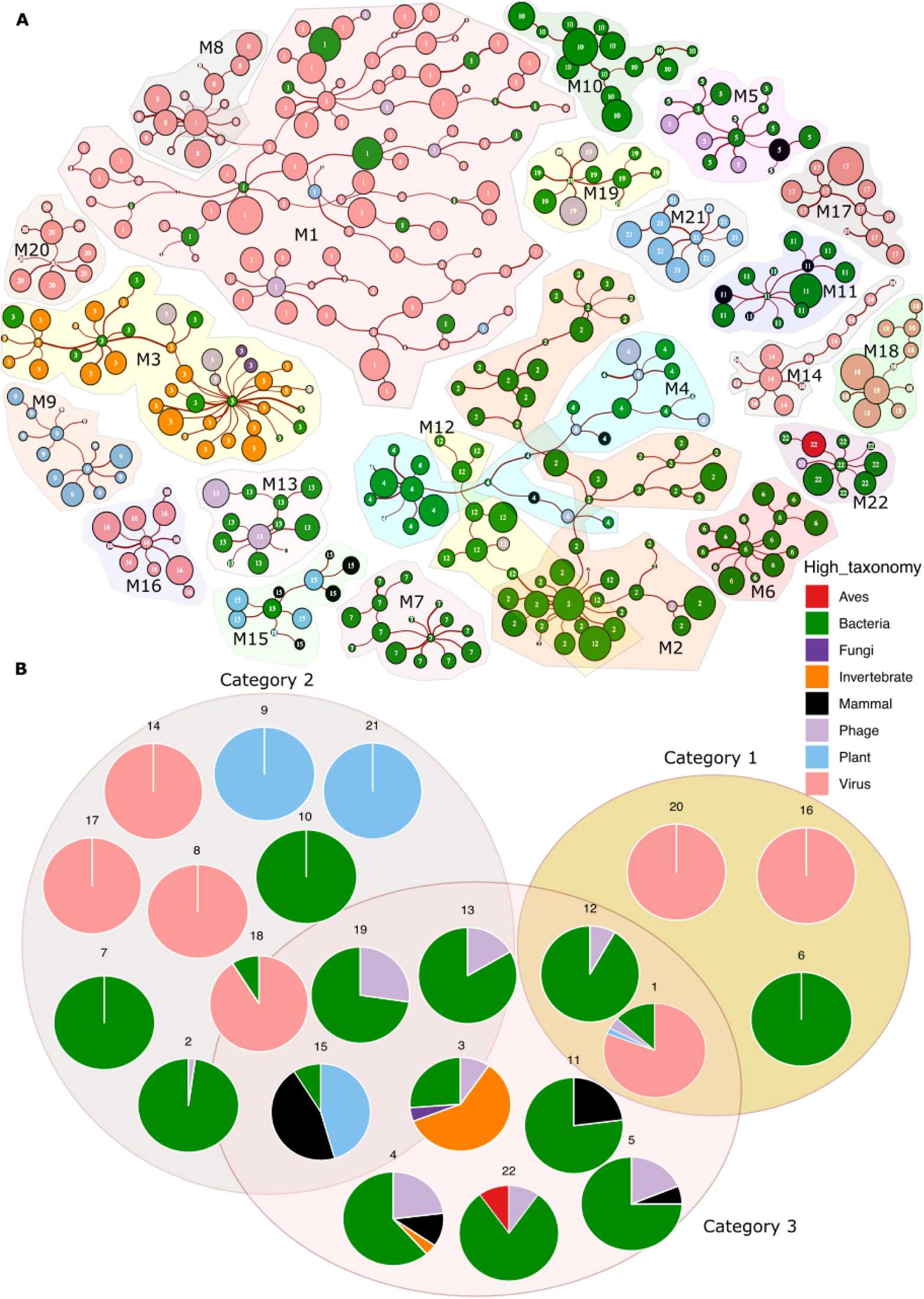
Co-occurrence network. Weighted gene network analysis identified 22 different antibody-bound peptide co-occurrent modules with at least 10 members. **A.** A minimum spanning tree was used to create the network of peptides belonging to one of the 22 modules. Nodes represent peptides, and node size is proportional to the peptide prevalence. Edges bind nodes with at least 0.3 Pearson correlation (between binary profiles). Colors represent different taxonomic sources of the peptide. Shades group modules and are labeled “M + module number”. **B.** Pie charts showing the taxonomic relative composition of each module. Pie charts are grouped in three categories. At right, category (1) indicates modules composed of different peptides from the same species. At left, category (2) indicates modules composed of structurally related peptides. At bottom is category (3) in which a mix of unrelated peptides from different organisms are seen. Category (3) may overlap with modules where the majority of peptides belong to category (1) or (2).

Next, we studied whether genetically related individuals or those living in similar environments (co-housing) would show more similarity in antibody-bound peptide enrichment compared to unrelated individuals. To explore this, we used 26 family trios from the LLD population (Genome of the Netherlands Consortium, 2014) (note that most offspring are unlikely to currently cohouse with their parents as mean age = 37±10.1 years old). Mother–offspring, father–offspring and father–mother antibody-bound peptide distances were significantly lower than those between unrelated individuals (p < 5×10^−4^, p = 0.013 and p < 5×10^−4^, respectively; 2,000 label permutations). However, no significant differences were found between family members although father–offspring pairs were, on average, more distant [**Figure 1D**]. The role of common environment in shaping antibody repertoires is supported by the decreased father– mother distance, while offspring associations could indicate an important role for environment during early life, a common lifestyle, the effect of genetics, or all to some degree.

### Co-occurrence of peptides identifies multiple epitopes for the same antigen, antibody cross-reactivity in related structures and co-occurrence of antibodies against unrelated structures

To understand the relation between antibody-bound peptides, we computed their correlation and built a network using weighted gene co-expression network analysis by computing correlation coefficients from the binary profile of all selected peptides without missing values. all, 435 peptides could be assigned to 22 modules of at least 10 highly correlated peptides [**Figure 2**] [**Supplementary Table 1.1**]. Assessing antibody-bound peptides within each of the modules (denoted by the number of peptides per module, 1 to 22) and the sequence similarity between them allowed us to identify three main types of modules: (1) modules driven by antigens from the same biological source, (2) modules driven by antigens of similar peptide sequence and (3) modules that include peptides that are not taxonomically or structurally related, but do correlate strongly with each other [**Supplementary Table 1.3**].

We observed five category (1) modules [**Figure 2**]. For example, module 16 was composed of two different Epstein-Barr virus (EBV) proteins, including capsid protein VP26 and nuclear antigen 1 (EBNA-1); module 20 was composed of high-identity peptides belonging to different strains of Influenza B viruses and module 1 is mainly driven by CMV peptides, while also including some EBV and other peptides. All modules are described in **Supplementary Table 1.3**.

Category (2) modules, driven by similar sequences in different peptides, highlight the cross-reactivity of antibody response [**Figure 2**]. For example, module 21 is composed of plant thionins, small cytotoxic plant compounds produced by many species, but here mainly derived from common wheat (*Triticum aestivum*), barley (*Hordeum vulgare*) and rye (*Secale cereale*). Module 9 contained related antigens from wheat, Asian rice (*Oryza sativa*), rye, barley and grass (*Setaria italica*) that represent plant granule–bound starch synthase peptides. Modules 14, 17 and 18 were characterized by antibody-bound peptides representing genome polyproteins from a series of viruses, including Enterovirus A71, B and C; Rhinovirus B and serotype 2; Coxsackievirus (type A9) and Poliovirus. Module 3 was dominated by allergen peptides, including antigens involved in common insect and seafood allergies, e.g. *Artemia franciscana* (shrimp), *Octopus vulgaris* (octopus), *Blattella germanica* (German cockroach), *Dermatophagoides farinae* (house dust mite), *Portunus trituberculatus* (gazami crab), *Bombus hypocrita* (bumble bee) and *Ctenocephalides felis* (cat flea).

Examples from category (3), where no structural or taxonomic relation is seen, are harder to interpret [**Figure 2**]. While some members in this category have a majority of peptides belonging to category (1) or (2), others do not show major structural relations and are mainly composed of bacterial peptides or bacterial and autoimmune peptides clustering together.

### Peptide enrichment is associated to HLA, FUT2 and IGHV genetic regions

Our observation that both common environments and genetic relations within families affect the antibody-bound peptide repertoire [**Figure 1E**] made us wonder about the specific drivers of repertoire variability. Genetics are known to influence antibody repertoires (Grundbacher, 1974; Kalff and Hijmans, 1969; Rowe et al., 1968; Venkataraman et al., 2021), but the exact contribution of genetic and environmental factors to bacterial and, especially, commensal gut microbiota immune-reactivity is incompletely characterized.

We estimated the proportion of antibody-bound peptide presence/absence variability accounted for by common genetic variation, i.e. its heritability (H^2^), using common genetic variants in 1,255 unrelated individuals. We saw an overall moderate genetic contribution to the variability of antibody-bound peptides enrichment (mean H^2^ = 0.1, median = 0.06, min = 0, max = 0.96) [**Supplementary Table 1.1**]. A total of 35/2,814 antibody-bound peptides showed very high heritability (H^2^ ≥ 0.5), while a substantial number (597/2,814) had a relatively high heritability (H^2^ ≥ 0.2). Using the highly heritable antibody-bound peptides (H^2^ ≥ 0.5), we then computed genetic correlations in order to determine similar genetic signals across antibody-bound peptide presence. We found a correlation of 0.47 between the matrices of presence/absence and genetic correlations (Mantel test, p < 1×10^−04^, 9,999 permutations) [**Supplementary Fig 1C**]. We also observed hubs of highly genetically correlated groups of peptides in which the genetic signatures are more correlated than antibody-bound peptide presence itself [**Supplementary Fig 1C**]. This indicates the existence of a common genetic architecture explaining the presence of antibody-bound peptides.

Next, we set out to uncover specific loci contributing to the observed heritability. We ran a genome-wide association study (GWAS) on 1,3640,125 genotyped and imputed SNPs in 2,815 peptides. To reduce the false discovery rate (FDR) and increase the power of the analysis, we meta-analyzed the results of our LLD GWAS with those of a dataset that used the same PhIP-Seq libraries in the context of inflammatory bowel disease (IBD) (490 participants), bringing us up to a total of 1,745 individuals (Bourgonje et al., in prep.)[**Supplementary Table 2.2**]. At study-wide significance threshold (p < 5.67×10^−11^), we identified three genetic loci associated to 149 antibody-bound peptides. These were located in chromosome 6 (Human leukocyte antigen (*HLA*) locus), chromosome 14 (Immunoglobulin heavy chain variable (*IGHV*) region) and chromosome 19 (fucosyltransferase 2 (*FUT2*) gene) [**Figure 3A**].

**Figure 3.**
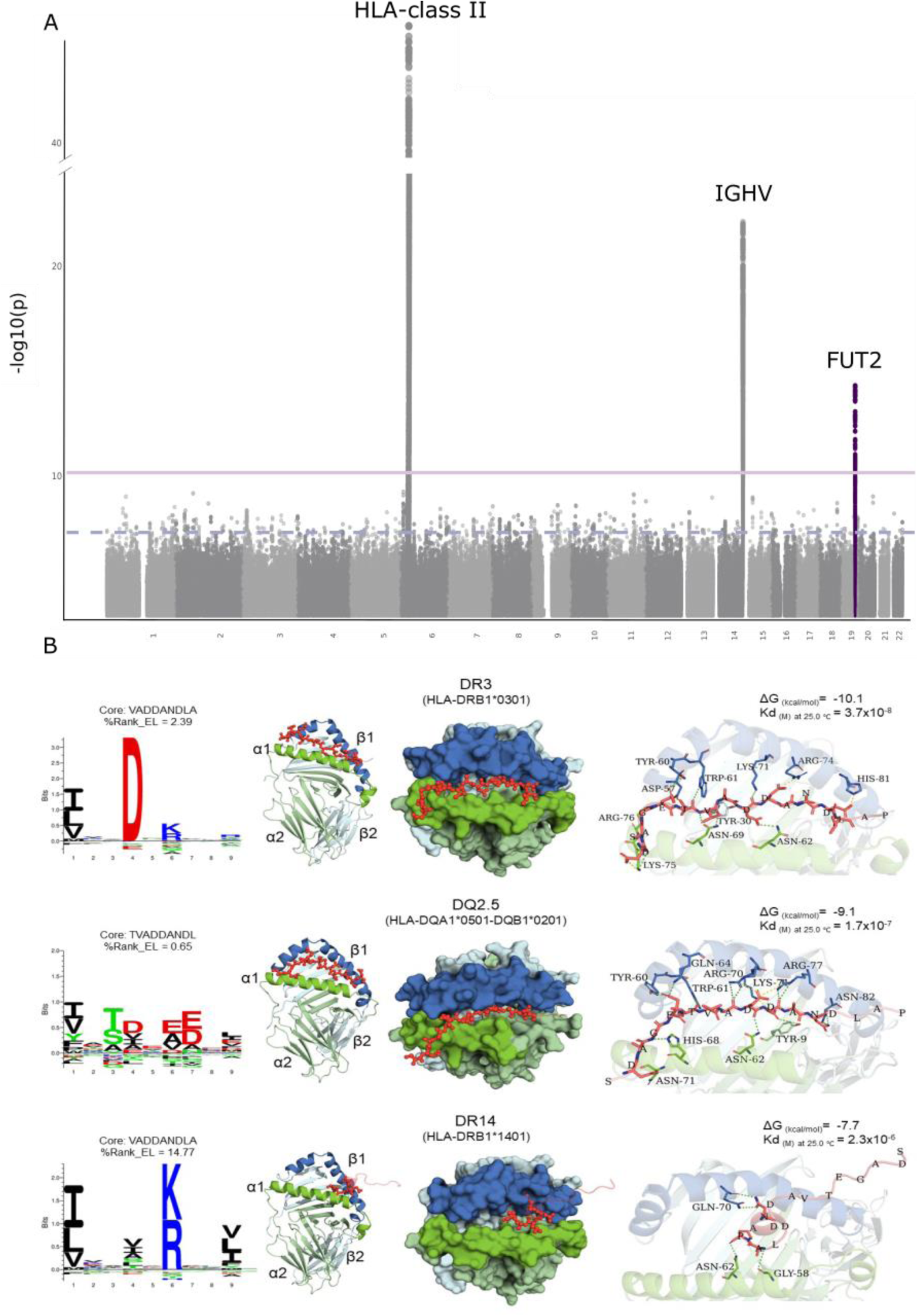
The genetic contribution to antibody-bound peptide variability. **A.** Manhattan plot from genome-wide association study of 2,798 antibody-bound peptides. Genome-wide association threshold (5×10^−8^, blue) and study-wide significance (7×10^−11^, red) are shown as horizontal lines. Labels indicate the three major loci identified. Colored dots represent a recessive model. Gray dots represent additive models. **B.** Peptide motif deconvolution maps of DR3, DQ2.5 and DR14 (amino acids code: negatively charged = red, positively charged = blue, polar uncharged = green, hydrophobic = black) compared with the Streptococcus agalactiae C5a peptidase peptide core and percentage of elution score (%Rank_EL: strong binding ≤ 2.0, weak binding 2.0–10.0, no binding > 10) predicted by NetMHCIIpan-4.0 (Reynisson et al., 2020a). Predicted binding mode, polar molecular interactions (dashes, hydrogen bonds: green, salt bridges: yellow), binding energy and dissociation constant (Kd) of the Streptococcus agalactiae C5a peptidase peptide core (red cartoon and sticks) into HLA-II receptors (chain A in green and chain B in blue).

The strongest genetic signal belonged to the HLA-class II region in chromosome 6, where we found 130 peptides associated with 134 different leading SNPs. Most of the associated peptides belonged to *Streptococcus* and *Staphylococcus* species, but we also found several peptides belonging to human viruses (adenoviruses or herpesviruses) and phages and to allergens (ovomucoid, barley, casein and wheat, amongst others) and gut microbiota. To dig into this genomic region, we conducted a specific imputation of HLA SNPs, indels, amino acids and gene isoforms and performed an association analysis with all peptides (see Methods) [**Supplementary Table 2.3**]. This analysis substantially increased the number of associated peptides and the strength of associations. We discovered that a large number of peptides (530/2,813) had at least one significant (p < 1×10^−6^, after correction for number of independent tests, see Methods) association with HLA variants (amino acids, insertions, SNPs or genes). At HLA gene level, we identified 1,267 significant peptide–gene associations to 276 different peptides. Most of those associations (and the strongest) belonged to allelic variants of HLA-II (1,139 associations to 271 different peptides) in comparison to variants of HLA-I (128 associations to 41 different peptides). Within the HLA-II variations, most associations were observed for various alleles in DQ and DR beta chain genes.

To determine whether these associations are due to the capacity of a specific HLA complex to present the peptide, we performed computational modeling of the HLA–peptide complex using some of our top associations. Here we identified that the predicted residues that are recognized from the peptide by a specific HLA complex (Reynisson et al., 2020a) can form stable structures with their associated HLA complexes. For instance, the streptococcal C5a peptidase (*TPSDAGETVADDANDLAPQAPAKTADTPATSKATIRDLNDPSQVKTLQEKAGKGAGTVVAVID A)* is highly associated with DRB1*0301 (always bound to the alpha chain DRA*01, DR3 haplotype) (Odds ratio (OR) = 3.78, p = 1.65×10^−31^) and with DQB1*0201 (OR = 3.75, p = 5.16×10^−31^) and the alpha chain DQA1*0501 (OR = 1.91, p = 4.80×10^−13^), which together form the haplotype DQ2.5 that is highly linked to DR3. The predicted core recognized by the HLA complex was nearly identical for both DR3 and DQ2.5 (*VADDANDL*) and is very similar to the amino acid composition identified from HLA ligand elution experiments (Reynisson et al., 2020b). This reveals a close relationship in the composition of the streptococcal C5a peptidase peptide sequence in comparison to chemically synthesized peptides tested for DR3 and DQ2.5. Additionally, we employed the predicted binding metric (percentage of elution score - %Rank_EL, Methods) to assess the binding of the core peptide to the selected alleles. %Rank_EL is calculated as the percentile of the predicted binding affinity compared to the distribution of affinities calculated on a set of random natural peptides (%Rank_EL; strong binding: ≤ 2.0, weak binding: 2.0-10.0, no binding: > 10). This analysis found a favorable binding prediction of the core to DR3 and DQ2.5 complexes, with a higher binding for DQ2.5 (2.39 and 0.65, respectively). We further compared the binding prediction for this epitope to a non-associated negative control (DR14) that was predicted to be non-binding (%Rank_EL 14.77).

Structural modeling and analysis of the binding mode of the peptide revealed a favorable binding energy with both DQ2.5 and DR3 (−9.1 and −10.1 Kcal/mol, respectively) compared to the non-associated structure DR14 (−7.7 Kcal/mol) [**Figure 3B**]. Additionally, the computed dissociation constant (Kd) showed an order of magnitude less affinity for the non-associated allele (2.3×10^−6^ M) compared to DR3 (3.7×10^−8^ M) and DQ2.5 (1.7×10^−7^ M). As a result, the peptide core exhibited similar behavior and key stabilizing polar interactions when binding into the binding sites of DR3 and DQ2.5. For example, the hydrogen bonds occurring between the Tyrosine 60 (Tyr60) and Tryptophan 61 (Trp61) present in the beta chain of both DR3 and DQ2.5 interact with Glutamic acid (Glu) and Threonine (Thr) in the peptide core. By contrast, although we could model the peptide binding into the negative control DR14, the majority of the peptide’s amino acids are located outside of the binding site and in the opposite direction compared to DR3 and DQ2 [**Figure 3B**].

In addition, we selected two other highly associated HLA–peptide complexes to explore in detail: (1) the combination of the peptide *Lactococcus* phage (YP_009222335.1 hypothetical protein LfeInf_097) with the DR15 haplotype (DRB1*0301), which showed the strongest study-wide association (OR = 13.3, p = 1.44×10^−47^) [**Supplementary Figure 2A**], and (2) a combination of a peptide from the *Human mastadenovirus* minor core protein with the associated DR4-DQ8 haplotype (encoded by the DRB1*0401 and DQA1*0301-DQB1*0302 genes) (DRB1*0401, OR = 5.69, p = 4.45×10^−15^; DQA1*0301, OR = 2.55, p = 2.12×10^−18^; DQB1*0302, OR = 3.14, p = 4.17×10^−20^) [**Supplementary Figure 2B**]. We observed a positive identification of the peptide core matching known deconvolution motifs, as well as a favorable binding prediction for the *Lactococcus* phage peptide to DR15 and for the *Human mastadenovirus* peptide to DR4-DQ8 haplotypes. Similarly, the binding mode modeling of the peptide cores to the HLA-II complexes resulted in energetically favorable binding energy calculations and Kd in the nanomolar range (*Lactococcus* phage–DR15, 1.6×10^−7^ M; *Human mastadenovirus*–DR4/DQ8, 1.2×10^−7^M and 1.3×10^−7^M, respectively). These results suggest that the identified HLA–peptide associations point to biologically relevant processes in which a specific HLA complex can preferentially bind and display the specific peptide sequence.

A second study-wide significant signal in our GWAS pointed to the *IGHV* region in chromosome 14 that encodes the immunoglobulin heavy chain variable domain. Here, we found 16 associated peptides in 11 leading loci within the region. The majority of SNPs (11/16) were located in non-coding regions around the *IGHV* gene, whereas *Ovis aries* casein protein (representing the primary sheep’s milk allergy food allergen) was associated with a missense variant that changes Glycine, a non-polar amino acid, for Arginine, a positively charged amino acid. Next to the *Ovis aries* casein peptide, the top peptides associated to this region are bacteria-related (*Bacteroides uniformis*, *Blautia producta* and *Lactobacillus plantarum*) or viral (*Influenza A*, *Lactobacillus phage* and *Norwalk virus*). The strongest association was observed in *Lactobacillus plantarum* (aggregation promoting factor) and *Lactobacillus* phage (endolysin).

We found a third study-wide significant signal in the *FUT2* gene in chromosome 19. This gene status controls the secretion or non-secretion (homozygous for loss of function) of the H-antigen, an oligosaccharide. Thus, we subsequently ran the analysis in a dominant/recessive model to increase power and detected three study-wide significant peptides, all of which originally belonged to Norwalk virus polyproteins and were negatively associated with the same leading variant, *rs2251034* (A>G,3’ UTR). This variant is in high linkage with an early-stop variant in *FUT2* that is known to stop the secretion of the H-antigen, rs601338 (A>G, R*^2^* = 0.85, 1000G, CEU population). FUT2 secretor status has been previously associated with multiple phenotypes, including infection susceptibility (Tian et al., 2017), gut microbiome (Kurilshikov et al., 2021; Lopera-Maya et al., 2020), human milk oligosaccharides (Williams et al., 2021) and cardiovascular traits (Zhernakova et al., 2018). Our finding supports the previously reported association between Norwalk virus susceptibility and FUT2 secretor status (Lindesmith et al., 2003), since this virus requires the H type-1 oligosaccharide ligand for successful attachment in the cell surface.

### Phenotypic and environmental effects on antibody-bound peptide enrichment

More than 200,000 bacterial antigens, including proteins originating from pathogenic, probiotic, and commensal gut microbiota species, were included in the peptide libraries. We therefore explored the relations between gut microbiome composition, analyzed by metagenomics sequencing, and presence of antibody responses. To increase the power of the study, we performed taxonomic abundance–peptide associations in 1,051 LLD participants and then ran the meta-analysis including 137 IBD participants (Bourgonje et al., in prep). Neither the cohort-specific analysis nor the meta-analysis strongly supported taxonomy metagenomic association with antibody-bound peptides (minimum FDR 0.52) [**Supplementary Table 2.4**]. These results are also in line with previous observations (Vogl et al., 2021).

To uncover specific effects of lifestyle and environmental factors in the antibody-bound peptide profile, we associated 84 available phenotypes [**Supplementary Table 1.2**] with the presence/absence of antibody-bound peptide profiles in 1,437 LLD participants. Here, we uncovered 837 strongly supported associations between the presence of antibody-bound peptides lifestyle and environmental factors (FDR < 0.05), covering 544 peptides and 48 different phenotypes [**Figure 4A**] [**Supplementary Table 2.5**]. Phenotypic factors that were associated (after age, gender and sequencing plate correction) with most antibody-bound peptides included age (386 associations), lymphocyte counts (101 associations, both absolute counts and cell proportions), neutrophil counts (86 associations, absolute counts and cell proportions), smoking (84 associations, both former and current smoking), sex (43 associations), allergies (35 associations, including any, pollen, dust or animals), autoantibodies (40 associations) and blood cholesterol levels (13 associations, both total cholesterol and LDL-cholesterol).

**Figure 4.**
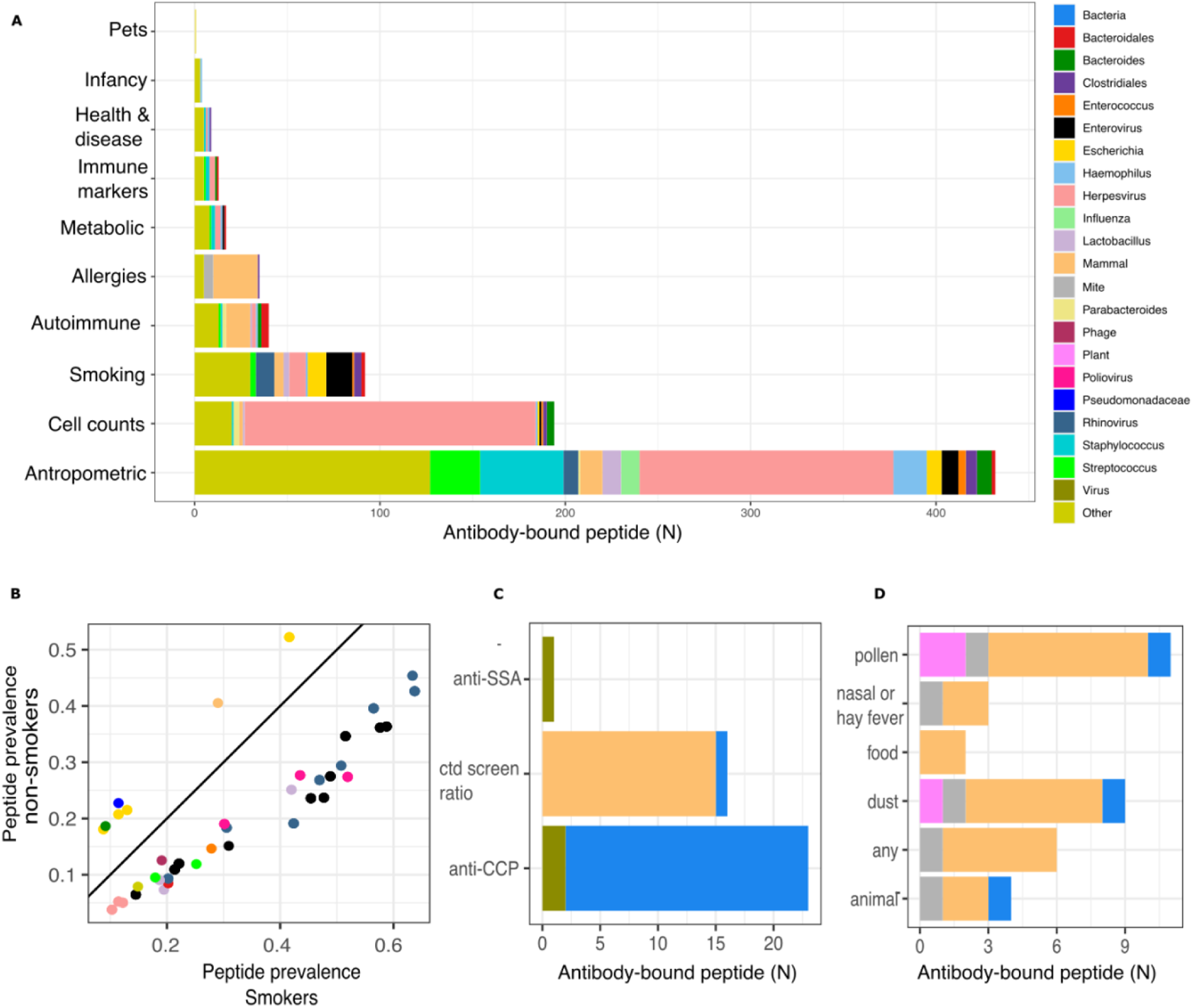
Phenotype-antibody-bound peptide associations. **A.** Bar plot displaying the number of associations per phenotype (FDR < 0.05). Phenotypes are grouped in categories. Peptides associated with > 5 phenotypes are grouped. Peptides associated with < 5 phenotypes are labeled ‘Other’. **B.** Smoking-linked antibody-bound peptide prevalence. X-axis shows prevalence of peptides in smokers. Y-axis shows the prevalence in non-smokers. Colors of dots depict peptide taxonomy. **C,D.** Autoimmune- and allergy-specific association counts of antibody-bound peptides, per category. Bacterial peptides are binned as “Bacteria”. Viral peptides are binned as “Virus”. Auto-antigens or antigens to casein are binned as “Mammal”. Plant peptides are binned as “Plant”. Anti-SSA: anti–Sjögren’s-syndrome-related antigen A autoantibodies. Anti-CTD: anti-connective tissue diseases screening ratio. Anti-CCP: anti-cyclic citrullinated peptide.

Of the 386 significant associations with age, 199 were positive and 187 were negative. Older age was associated with a higher prevalence of antibody-bound peptides from several herpes viruses (including CMV, EBV and Herpes simplex virus (HSV) 1 and 2), *Streptococcus* bacteria (in particular *S. pyogenes* and *S. dysgalactiae*) and several pathogenic bacteria (including *Shigella flexneri*, *Yersinia enterocolitica*, *Campylobacter* genus and *Helicobacter pylori*). Younger individuals had higher frequencies of antibody-bound peptides related to particular viruses (including human rhinovirus serotype 2, influenza A virus and enteroviruses) and bacteria, mainly *Streptococcus pneumoniae*, *Staphylococcus aureus*, *Mycoplasma pneumoniae*, *Haemophilus influenzae* and *Escherichia coli* (particularly antigens from the type III secretion system (T3SS) of serotype O157:H7). Younger individuals also showed more frequent antibody responses against alpha S1 casein proteins.

Sex demonstrated 43 significant enrichments (24 for males, 19 for females). Females exhibited more frequent antibody-bound peptides from *Lactobacillus acidophilus* and *Lactobacillus johnsonii*, both known inhabitants of the vaginal microbiome (Davoren et al., 2019; Integrative HMP (iHMP) Research Network Consortium, 2019). Antibody-bound peptide responses were particularly directed against *Lactobacillus* surface proteins, including S-layer proteins (SLPs, e.g. SIpA and SIpX proteins) and the peptidoglycan lysozyme N-acetylmuramidase, reproducing previous findings in (Vogl et al., 2021). Females also demonstrated increased enrichment of EBV and CMV peptides. Males showed higher prevalence of antibody-bound peptides from *Haemophilus influenzae* bacteria (e.g. serotype Rd KW20 or strain 3179B), also as previously described (Angkeow et al., 2021; Kurtti et al., 1997), and of several peptides derived from *Streptococcus, Staphylococcus, Bacteroides* and alphaherpesviruses (including HSV-1 and varicella zoster virus).

Associations between antibody-bound peptides and laboratory cell counts included both cell proportions and absolute cell quantifications, which appeared to be largely driven by antibody-bound peptides from CMV. Lymphocyte counts showed almost exclusively positive associations with CMV, but also some to EBV, whereas the same antibody-bound peptides demonstrated many inverse associations with neutrophil counts.

Smoking associations included associations to current smoking status (41) [**Figure 4B**], ever smoking for at least a year (43) and parental smoking (7). Most associations were related with higher prevalence of peptides belonging to enteroviruses, both rhinovirus and poliovirus. The relationship between smoking and rhinovirus infection has been previously described (Cohen et al., 1993), and thus associations to other viral peptides belonging to enteroviruses could be due to cross-reactivity to homologous proteins. We also observed a consistently higher seroprevalence of EBV in smokers, which might be reactivated by smoking, as shown by an *in vitro* model (Xu et al., 2012). In addition, there were increased antibody responses against miscellaneous respiratory pathogens, including several *Streptococcus* spp. Interestingly, flagellin antibody–bound peptides (*Roseburia*, *Lachnospiraceae*, *Eubacterium* and *Clostridiales*) show a lower prevalence in smokers, as do *Escherichia* virulence factors [**Figure 3B**].

We used serological information about the presence of autoantibodies to identify bacterial and allergen peptides linked to the presence of these autoimmune antibodies [**Figure 4C**]. Anti-cyclic citrullinated peptide (anti-CCP) antibody levels, a marker for rheumatoid arthritis, were positively associated with 23 antibody-bound peptides, including peptides derived from *Bacteroides*, *Parabacteroides*, *Prevotella* and *Porphyromonas gingivalis* bacteria. These findings correspond well with bacterial genera that are known to be altered in the microbiome of patients with anti-CCP-positive rheumatoid arthritis (Bodkhe et al., 2019). On the other hand, the connective tissue disease (CTD) screen panel, in which total reactivity to a mixture of antigens associated with several autoimmune diseases is measured, was almost exclusively associated with increased antibody-bound peptide frequencies of alpha-S1-casein or kappa casein belonging to *Bos taurus* (cow), *Ovis aries* (sheep), *Bubalus bubalis* (buffalo) and *Capra hircus* (goat). Indeed, several autoimmune diseases such as celiac disease, juvenile idiopathic arthritis and Ehlers-Danlos syndrome have been associated with mucosal reactivity against milk allergy, where the casein protein seems to be a regulator of the inflammatory response (Cutts et al., 2012; Kristjánsson et al., 2007). Anti-Sjögren’s-syndrome-related antigen A antibodies (anti-SS-A/anti-Ro), which are typical anti-nuclear antibodies associated to autoimmunity, were positively associated with an antibody-bound peptide representing thymidine kinase of EBV. Interestingly, this association has previously been described in the context of Sjögren’s syndrome, in which anti-SS-A autoantibodies and higher frequencies of serological EBV reactivation (Fox et al., 1991) are more frequently observed.

The strongest association to total cholesterol levels was with an antibody-bound peptide of *Haemophilus parainfluenzae* strain T3T1. Other bacterial peptides are also enriched with higher cholesterol labels, including *Streptococcus* or *Pseudomonadaceae*. We also observed an enrichment of viral peptides, such as rubeola, *Pneumoviridae*, HSV and EBV. Many intracellular pathogens are known to use cholesterol drafts to successfully infect cells and to impair the regular cholesterol metabolism and the immune system (Sviridov and Bukrinsky, 2014). We observed three associations between body-mass index (BMI) and antibody-bound peptides, all of which represented glycoprotein D of human alphaherpesviruses (HSV-1/HSV-2). Indeed, obesity has previously been associated with a higher prevalence of herpesvirus infections, in particular HSV-1, by promoting human adipogenesis (Hasan et al., 2021).

Finally, participant’s having any allergy (44.5% of participants) showed associations with six different antibody-bound peptides [**Figure 4D**]. Using more-detailed questionnaires with information about different allergies such as dust, pollen, food and others [**Supplementary Table 1.3**], we identified 13 different peptides associated with at least one phenotype. As expected, the strongest association was observed for dust allergy, showing associations with antibody-bound peptides from the house dust mite *Dermatophagoides pteronyssinus* (FDR = 3.84×10^−5^). In addition, the most common associations were observed between casein proteins derived from cow, sheep and buffalo milk, which were linked not only with food allergies but with almost all allergy types. Wheat allergens were linked with self-reported dust and pollen allergies. Interestingly, we identified a couple of associations with influenza (higher prevalence with pollen allergy), bacterial flagellin associations with animal allergies and *Shigella flexneri* with dust allergy. Previous analyses have linked dust mite with bacterial sensitization, although not for these specific lineages (Dzoro et al., 2018). Importantly, several of these significant associations represent linkage between common aeroallergens (e.g. pollen and dust) and food allergy (e.g. *Triticum aestivum* [wheat] and casein), recapitulating the frequent co-occurrence of allergen cross-reactivity (Popescu, 2015).

## Discussion

In this project, we aimed to characterize the antibody repertoire in the blood of a Dutch population and reveal which factors contribute to its variation. In particular, the factors that contribute to the generation of antibodies against microbiota and different allergens remain elusive. Here, we combined phenotypic and genetic information together with the immune-interrogation of 2,815 common peptides from microbes, viruses, allergens and self-peptides to study this variability. Using population, family and longitudinal samples, we identified the antibody profile in the general population, assessed the stability of antibodies after 4 years and investigated the effect of genetic and environmental factors on individual immune profiles.

The relation between genetics and antibody repertoire has been extensively described (Grundbacher, 1974; Kalff and Hijmans, 1969; Rowe et al., 1968; Venkataraman et al., 2021) but has been limited to a relatively small number of antibodies until now. PhIP-Seq has recently enabled the investigation of the genetic contribution to antibody variability in a much broader scale, although it has mainly been investigated for viruses, toxins and virulence factors (Angkeow et al., 2021; Venkataraman et al., 2021) and not for other antigens such as allergens and gut microbiota–derived proteins. Here, we identified three genomic regions highly associated with the variability of antibody-bound peptide repertoires. As expected, we replicated the relation between HLA loci and antibody-bound peptide prevalence (Angkeow et al., 2021; Kachuri et al., 2020; Scepanovic et al., 2018; Venkataraman et al., 2021). Through imputation of HLA alleles, amino acids and structural variants, we also set out to uncover the specific HLA variations that allow the peptide to be displayed. Our structural simulations of the HLA alleles agree with the observed association patterns, supporting the hypothesis that the strong associations are due to HLA-display capabilities. For the first time, we report massive and specific HLA associations to more than 500 peptides at a high confidence level. This association data will be used in the future to further understand HLA–peptide interactions by modeling possible residue interactions. Our findings also support previous observations, such as the association of *FUT2* and Norwalk virus peptides (Lindesmith et al., 2003) that is explained by the attachment of the viral particle to the epithelia of *FUT2*-secretor cells (Marionneau et al., 2002). We also observed association in the *IGHV* locus that was not previously reported in relation to antibody profiles. This association is in a complex genetic region as several genes with multiple isoforms coexist in the genome that are hard to address with microarrays (Watson and Breden, 2012). In addition, we lack information about the rearrangements that this gene undergoes during B-cell maturation. Nevertheless, although we cannot directly interpret the relation between variation and peptide recognition, this is a genetic region that is expected to contribute to antibody-bound peptide variability. Interestingly, our study did not identify the previously reported association of the nucleoredoxin gene (*NXN*) with *S. pyogenes’* M3 Streptolysin O (*SLO*) protein (Angkeow et al., 2021), although we do find a weak positive association between rs4968063 and the prevalence of this antibody-bound peptide in the combined LLD and IBD cohort (p = 0.01).

In the present study, we observe a lack of concordance between meta-analyzed fecal microbial composition and PhIP-Seq-based epitope repertoires, which is in line with findings from studies using the exact same library of antigens in a healthy population-based Israeli cohort and in a disease cohort consisting of patients with IBD (Vogl et al., 2021; Bourgonje et al., in prep). The top associations do not present clear relationships between specific microbial taxa and antibody-bound peptides, which could be explained in various ways. First, this apparent lack of association might point to past events, such as microbial translocation, that may have triggered long-lasting immunity that was captured by PhIP-Seq profiling (Marchix et al., 2018), while the respective bacteria have been cleared from the gut. Second, there may have been a lack of resolution in the microbiome data. For example, some bacterial species commonly detected by metagenomics may have been accompanied by higher detection thresholds in PhIP-Seq, whereas highly immunogenic antigen peptides may not be frequently detected by metagenomics sequencing (Vogl et al., 2021). In addition, the use of fecal microbiota as a proxy for the gut microbiota limits the characterization of local immune– microbiota interactions. Profiling mucosa-attached microbiota rather than fecal microbiome could have improved the antibody-bacteria concordance as locally residing (mucosal) microbial communities may elicit stronger immune responses that may also depend on the anatomical location within the intestines (Christmann et al., 2015).

We also explored the relationship between peptide prevalence and various morphological, biochemical and lifestyle factors. Our observations reveal a number of interesting associations. For instance, EBV and CMV were associated with lymphocyte and neutrophil counts. These findings are in accordance with observations of absolute lymphocytosis and neutropenia that constitute characteristic laboratory findings in individuals affected by EBV (infectious mononucleosis) (Fisher, 1973; Hudnall et al., 2003) or CMV infections (Lima et al., 2006; Solana et al., 2012), which may translate into altered immune cell proportions on the longer-term. We also identified a series of associations of allergies and allergens. Allergies are normally triggered by the epitope interaction with IgE antibodies.

However, in this study, we mainly used IgG for immunoprecipitation since IgE are found in small amounts in serum and bind with relatively low affinity to the protein A/G coated magnetic beads employed for the immunoprecipitation. Previous studies have shown that allergens have the chance to bind both to IgG and IgE, although they might have different epitope preferences (Monaco et al., 2021). Thus, the allergen associations presented here should be interpreted with caution as they may differ from the classical pathway involved in allergy.

Using co-occurrence networks, we identified different peptide groups that normally belonged to the same taxa or orthologous structures in different taxa. However, the existence of modules with apparently unrelated peptides may indicate either interesting biological phenomenon or technical factors that we are not accounting for. For instance, *H. pylori* peptides were seen to occur with the prevalence of antibody-bound peptides for a couple of phages. The disruption of the gut barrier by this pathogen (Fukuda et al., 2001) could potentially explain the translocation of those phages to blood and the generation of mucosal and systemic immune responses. On the other hand, phenotypic associations also allow us to conjecture about observed cryptic peptide co-occurrence. For instance, CMV peptides were seen to co-occur with several bacterial and plant peptides. Most of those peptides were associated to the same phenotypes, mainly blood cell leukocyte and granulocyte counts, age and sex, meaning that the co-occurrence could be driven by those factors, or that those phenotypes may mediate their co-occurrence.

Widespread antibody screenings will be of great importance for the immunology field. Large longitudinal studies will enable us to go from association to causality, for instance uncovering factors that influence the development of autoimmune diseases (Elkon and Casali, 2008) or common allergies (Kearney et al., 2015). Studies like this will enable the development of personalized treatments, e.g. through vaccination strategies (Cotugno et al., 2019).

## Study limitations

PhIP-Seq is currently limited to linear epitopes and lacks post-translational modification information, and thus new technologies or improvements of the current method (e.g. as in (Román-Meléndez et al., 2021)) are still to be developed. Similarly, the nature of the assay will also miss tridimensional structure information from the antigens that might be recognized by the antibodies. In addition to these technological issues, our relatively small sample size for genetic studies hampers an accurate estimation of antibody-bound peptide heritability and genetic correlation. It is also important to acknowledge that the antibody-bound peptides we identified mainly correspond to circulating IgG and may overlook other types of immunoglobulins or immunoglobulins not in systemic circulation. Finally, due to the mostly cross-sectional nature of the experimental design, it is hard to draw causal links from the associations we present and further studies are needed to establish causality and dependence.

## Materials & Methods

**Lead contact:** Further information and requests for resources should be directed to the Lead Contact, Alexandra Zhernakova.

**Material availability:** List of antibody-bound peptides enriched in LLD participants will be made available upon publication.

**Data code and availability:** https://github.com/GRONINGEN-MICROBIOME-CENTRE/Phip-Seq_LLD-IBD

### 1. Cohort information

Lifelines is a multi-disciplinary prospective population-based cohort study examining, in a unique three-generation design, the health and health-related behaviors of 167,729 individuals living in the North of the Netherlands. It employs a broad range of investigative procedures to assess the biomedical, socio-demographic, behavioral, physical and psychological factors that contribute to the health and disease of the general population, with a special focus on multi-morbidity and complex genetics (Scholtens et al., 2015). We collected data from the subcohort LLD (Tigchelaar et al., 2015) (58% female, mean age 45.04 years, mean BMI 25.26, 12% obese participants with BMI > 30). Approval from institutional ethics review is available under reference number M12.113965. In this study, we used a subset of LLD (n = 1,437, 57% female, mean age 44.5 years) with available information including anthropometrics, blood parameters and self-assessed questionnaires about health and lifestyle. The autoantibody panels for anti-CCP and CTD-ratio and anti-SSA were originally described in (Lambers et al., 2021) and (van Zanten et al., 2017), respectively.

The 1000IBD cohort is a large, prospective observational cohort study based in Groningen, the Netherlands, aiming to biologically and clinically characterize patients with IBD who are included at the outpatient IBD clinic of the University Medical Center Groningen (UMCG) (Imhann et al., 2019). Detailed phenotypic data and multi-omics profiles have been generated for over 1,000 included patients with IBD, enrolled from 2,007 onwards. Antibody-bound peptide repertoires (PhIP-Seq profiles) were generated for 497 patients included in the 1000IBD cohort (median age 39 years, 63% females, median BMI 24.7 kg/m^2^), of which 256 patients were diagnosed with Crohn’s disease, 207 with ulcerative colitis and 34 with an undetermined type of IBD (IBD-U). Ethical approval for participation in the 1000IBD cohort has been granted by the Institutional Review Board of the UMCG (in Dutch: “Medisch Ethische Toetsingscommissie”, METc) under registration number 2008/338 and the study has been conducted in accordance with the principles of the Declaration of Helsinki (2,013). Patients provided written informed consent for their participation in the study. Further details on the subcohort of 1000IBD of which PhIP-Seq profiles were generated can be found elsewhere (Bourgonje et al, in prep).

### 2. PhIP-Seq library design, preparation, sequencing and processing

Library description can be found in (Vogl et al., 2021) (microbiota antigens) and (Leviatan et al, in prep) (allergen databases, complete IEDB, phages). The general PhIP-Seq protocol is described in (Larman et al., 2013) and was performed with minor modifications outlined by (Vogl et al., 2021). In short, PCR plates in contact with phage/antibody mixtures were blocked with bovine serum albumin (BSA) solution (concentration as described in (Vogl et al., 2021)). BSA was supplemented into phage-buffer mixtures for immunoprecipitations (IPs). Phage wash buffer for IPs contained 0.1% (wt/vol) IPEGAL® CA 630 (Sigma-Aldrich cat. no.I3021). Phage and antibody amounts for IPs were used as optimized by (Vogl et al., 2021) at 3 µg of serum IgG antibodies (measured by ELISA) and phage library at 4,000-fold coverage of phages per library variant. As technical replicates of the same sample were in excellent agreement (average Pearson ρ = 0.96, (Vogl et al., 2021)), measurements were performed in single reactions. The libraries (Vogl et al., 2021) (230 nt, 244,000 variants) were mixed in a 2:1 ratio with the phage, immune and allergen library (200 nt, 100,000 variants) (S.L., manuscript in preparation).

Phage–antibody mixtures mixed with overhead mixing at 4°C. A 50%-50% mix of protein A and G magnetic beads (total 40 μl; Thermo Fisher Scientific, cat. nos. 10008D and 10009D, prepared according to the manufacturer’s recommendations) was added after overnight incubation and further rotated at 4°C for 4 h, then the beads were transferred to PCR plates and washed twice, as previously reported (Vogl et al., 2021). Therefore, a Tecan Freedom Evo liquid-handling robot with filter tips was used.

PCR amplifications (pooled Illumina amplicon sequencing) were run with Q5 polymerase (New England Biolabs, cat. no. M0493L) according to the manufacturer’s recommendations (primer pairs as outlined by (Vogl et al., 2021)).

#### Composition of the antigen library

This work uses two previously developed peptide libraries: a microbial library (Vogl et al., 2021) and an allergen library (Leviatan et al, in prep). The microbial library contains 244,000 peptide sequences from 28,668 different proteins, from which 27,837 proteins were derived from microbial antigens, while the rest are controls. This contains genes predicted from metagenome assembled genomes (147,061 peptides), known pathogenic bacterial species (61,250 peptides), bacteria known to be coated with antibodies (22,050 peptides), probiotic bacteria (14,700 peptides), virulence factors extracted from the virulence factor database (VFDB) (24,164 peptides) and controls (11,525 oligos). Antigens were selected giving priority to known immunogenic antigens and focusing on secreted, membrane and motility proteins. The second library contained 5,527 peptides from five different allergen databases (Leviatan et al, in prep), 31,436 peptides from the Immune Epitope Database (IEDB) (Vita et al., 2015) and approximately 40,000 phage peptides.

#### Peptide antibody-binding enrichment

Antibody-binding against peptide (seropositivity) was defined as described in (Vogl et al., 2021). In brief, for each sample, null distributions per input level (number of reads per clone without IP) are generated. A two-parameter generalized Poisson model is fit to the null distribution, and the P-value to obtain the coverage level after IP for a given clone is estimated. Model parameters were estimated for each null distribution using maximum likelihood or directly interpolated as described in (Larman et al., 2011). A strict Bonferroni cut-off at P_Bonferroni_ < 0.05 was then used to define seropositivity. A total of 175,242 peptides were seropositive in at least one participant.

### 3. Antibody-bound peptides exploratory analysis

Data analysis was performed in R v4.0.3 using the packages tidyverse, stats, vegan (Dixon, 2003), corrplot, igraph (Csardi et al., 2006), WGCNA (Langfelder and Horvath, 2008), readxl, pheatmap, cairo and patchwork.

#### Antibody-bound peptide selection

Peptides to be used in the analysis were selected based on two filters. We chose peptides that had a prevalence at least of 5% and below 95% in either 1000IBD or LLD (excluding follow-up samples). For antibody-bound peptides with identical sequence, we chose the most prevalent antibody-bound peptide, resulting in 2,815 selected antibody-bound peptides.

#### Principal component analysis

We used 2,815 peptides to compute a PCA. Eigenvalues were used to produce a scree plot and eigenvectors to identify top peptides contributing to the first components. A K-means algorithm (k = 2) was performed on the dimensionally reduced dataset (PC1 and 2) to label observed clusters. This analysis was reproduced after removal of the 90 peptides belonging to CMV.

#### Time and family distance analysis

322 LLD samples belonging to two different time points were used for a time consistency analysis. Jaccard distance was used as the dissimilarity metric between samples. P-value of longitudinal effect of mean distance was estimated by computing the P-value of the mean pairwise difference of longitudinal samples in a null distribution of mean distances of pairwise differences of 2,000 label swaps. Interrogation of factors that might affect the degree of change in longitudinal samples was performed using pairwise distances from longitudinal samples as dependent variable and age and sex as covariates in a linear model. Antibody-bound peptide consistency was computed by averaging the number of changes in the enrichment profile of a peptide among all samples with longitudinal data points. To check whether antibody-bound peptide enrichment changes seen in follow-up are due to a different reactivity of the plates used for baseline and follow-up samples, we ran a Wilcoxon test comparing the number of enriched antibody-bound peptide of participants profiles from plates with follow-up samples vs plates with no follow-up samples.

We then selected samples belonging to the same family (Genome of the Netherlands Consortium, 2014) with three members (26 families). We computed pairwise distances (Jaccard) between family members (father to offspring, mother to offspring and father to mother). For each of the comparisons, we estimated a P-value comparing the mean distance with a random distribution of means between 2,000 permuted labels.

#### Network analysis

We used a weighted gene co-expression network analysis (Langfelder and Horvath, 2008) in the context of antibody-bound peptide presence/absence to identify modules of peptide co-occurrence. We used all LLD samples (1,784) and the subset of selected peptides with no missing values (2,770) to build the network. The soft thresholding power was chosen by visually inspecting the model fit of powers from 1 to 20. It was decided to use a power of 7. A network was built using Pearson correlation between antibody’s presence/absence profiles, followed by hierarchical clustering. A cut-off height for merging of 0.5 was used and a minimal module size of 10 peptides was required for a module to be called. The peptide identity from the identified modules was checked and a sequence similarity analysis was run. Module *eigengenes* were extracted using WGCA. *Eigengenes* were correlated between modules. Strong module correlation was defined on the basis of achieving a P_Bonferroni_ < 0.05.

Peptides belonging to a module of at least 10 peptides were used to build a visual network graph (igraph). A maximum spanning tree algorithm was used to build the network.

To check if co-occurrence modules might be driven by batch effects (due to PhIP-Seq plate), we computed the prevalence of each peptide within a module. If a common batch effect was present in all peptides of a module, we would expect to see a significant batch effect adding variation to the mean prevalence within all modules (Null hypothesis, Prevalence ~ Peptide + Batch). If this batch effect was different per peptide, then the batch effect would show a significant interaction with the peptide (Alternative hypothesis, Prevalence ~ Peptide + Batch + Peptide*Batch). If the alternative hypothesis was true, the batch would have a different effect per peptide, and thus it is not the only explanation to observe high co-occurrence between antibody-bound peptides. We fitted the null and alternative hypothesis in two linear models, and computed a P-value for the peptide–batch integration by computing a likelihood ratio test between both models. All tested models showed a significant interaction effect, indicating that batch most likely has a different effect per peptide.

#### Peptide similarity

Sequence similarity between peptide groups of interest was estimated using Clustal Omega (Sievers et al., 2011). Clustal Omega uses this distance matrix to build guiding trees for the progressive multiple sequence alignment algorithm. This distance is internally calculated using the k-tuple method (Wilbur and Lipman, 1983).

### 4. Phenotype association analysis

Jaccard distances between all samples were used as the dependent variable in a PERMANOVA against, sex, age and PhIP-Seq plate in order to identify covariates of interest. To associate individual enrichment profiles to available phenotypes, we performed a logistic regression on the presence/absence of antibody-bound peptides using the phenotype of interest, PhIP-Seq plate, age and sex as covariates on 1,437 baseline participants. We controlled the FDR at 0.05 using the Benjamini-Hochberg procedure (Benjamini and Hochberg, 1995).

### 5. Genetic analyses

#### Genotyping and imputation

Genome-wide genotyping data was generated as described in (Tigchelaar et al., 2015). Genotype data processing is described in (Zhernakova et al., 2018). Briefly, microarray data were generated on CytoSNP and ImmunoSNP platforms and processed on the Michigan Imputation Server (Das et al., 2016). Haplotype phasing was carried using SHAPEIT and imputation using the HRC version R1 as reference (Consortium and the Haplotype Reference Consortium, 2016).

#### Genetic preprocessing

We used GenotypeHarmonizer (Deelen et al., 2014) for imputation (minimum posterior probability of 0.4), call rate (minimal call rate of 95% of samples), Hardy-Weinberg equilibrium (minimal P-value allowed of 1×10^−6^) and SNP ambiguity filtering. We then computed identity by descent among samples using PLINK v1.9 (Purcell et al., 2007) on linkage disequilibrium (LD)– pruned genotypes (window size 50 Kb, variance inflation threshold 5 and maximum R^2^ between variants 0.2). We estimated identity by descendant between all samples using PLINK and randomly selected a sample from the pairs with a PI_hat value > 0.2, which resulted in the removal of 14 samples from subsequent analysis (total of 1,255 available samples).

#### Heritability and genetic correlation

GCTA (Yang et al., 2011) was used to compute a genomic relationship matrix (GRM) using genotyped SNPs with a minor allele frequency (MAF) of at least 0.05. The GRM was used to estimate antibody-bound peptide heritability using a linear mixed model between unrelated individuals (GREML approach) (Yang et al., 2010) while controlling for age, sex and PhIP-Seq plate. Similarly, genetic correlations between peptides were estimated using GCTA (Lee et al., 2012).

#### Genome-wide association

For each of the available antibody-bound peptides, we conducted an association analysis between genotypes (MAF > 0.05) and presence/absence profile. PLINK v1.9 (Purcell et al., 2007) logistic mode was run while controlling for age and sex and using the genotype in an additive model. This analysis was reproduced in a recessive model between 49.1 and 49.3 Mb in chromosome 19.

#### Genetic meta-analysis

A second study using the same PhIP-Seq library panel and protocol has been conducted in an IBD cohort from the Netherlands (Imhann et al., 2019; Bourgonje et al. under prep). Genotyping information is available for this cohort and was previously described in (Hu et al., 2021). The same quality control steps and analysis methods have been used as described above, while the disease subtype (Crohn’s disease or ulcerative colitis) was also added as an extra covariate in the logistic regression.

Summary statistics from both the LLD and 1000IBD cohorts were meta-analyzed using METAL (Willer et al., 2010). We performed a *P*-value–based fixed-effects meta-analysis. A study-wide significance threshold was estimated by dividing the genome-wide significance threshold of 5×10^−8^ by the number of independent peptides included in the GWAS. The number of PCs needed to reach 90% of antibody-bound peptide repertoire variability in LLD was used as a number of independent tests (708), obtaining a study-wide threshold of 5.67×10^−11^. For each peptide’s summary statistics we extracted genome-wide significant associations (p<5×10^−8^) for clumping. We clumped variants in windows of 1,000 Kb if they had a minimal R^2^ (computed from LLD genotypes) of at least 0.1 using PLINK. Leading variants of each clump were then annotated using the Ensembl Variant Effect Predictor and the grCh37 human build (McLaren et al., 2016). LD between our identified leading variants and other publicly reported variants was estimated in the CEU population from the 1,000 genomes using the LDlink webtool (Alexander and Machiela, 2020; Machiela and Chanock, 2015).

#### HLA imputation and association

The chromosome 6 region with 25–34 Mb that contains the MHC genes was extracted. Imputation of the HLA region, including HLA alleles, polymorphic amino acids, SNP variants and indels, was then performed using SNP2HLA (v2) with the Type 1 Diabetes Genetics Consortium (T1DGC) reference panel (2,767 unrelated European descent individuals) HLA Reference Panel (Jia et al., 2013). Next, we combined both imputed and genotyped SNPs, HLA alleles and amino acid variants, resulting in a total of 8,926 variants. Variants with MAF < 0.05 and imputation quality score (INFO) < 0.5 were removed before association.

HLA to peptide association was performed using linear models in 1,175 participants, while controlling for age, sex, PhIP-Seq plate and disease subtypes (Crohn’s disease/ulcerative colitis, only specific to IBD cohort). Summary statistics from both datasets were further meta-analyzed using a fixed-effects model in PLINK v1.9. The statistical significance threshold was determined by dividing the usual P-value 0.05 threshold level by the number of independent features tested (66 PCs were needed to reach 90% of HLA feature variability in LLD, while 708 PCs were needed to capture 90% of the peptide variability, resulting in 46,728 independent tests), resulting in a threshold of 1×10^−6^. FDR was estimated using the Benjamini-Hochberg method (Benjamini and Hochberg, 1995).

#### Modeling of peptide presentation in HLA complexes

To explore whether HLA–peptide associations potentially point to HLA-II ability to display a specific peptide, we performed computational modeling of the complex–peptide interaction.

The protein sequences of DR3, DR4, DR14, DR15 and DQ2 were obtained from the IPD-IMGT/HLA database (Robinson et al., 2020) and aligned against the entire Protein Data Bank database using pBLAST. Protein structures displaying a 100% of amino acid identity with the HLA-II database sequences were chosen to build the peptide binding modes. Those structures correspond to the HLA complexes DR3:7N19, DR4:1D5M, DR14:6ATF, DR15:1YMM, DQ2:6PX6 and DQ8:2NNA. Proteins other than HLA-II, water molecules and heteroatoms were removed from the structures prior to modeling. The NetMHCIIpan-4.0 (Reynisson et al., 2020a) server was then used to predict peptide binding to the corresponding associated HLA alleles: DRB1*1501 for *Lactobacillus phage* LfeInf; DRB1*0301, DQA1*0501-DQB1*0201 and DRB1*1401 for *Streptococcus agalactiae* C5a peptidase; and DRB1*0401 and DQA1*03-DQB1*0302 for Human mastadenovirus minor core protein. The DRB1*1401 for *Streptococcus agalactiae* C5a peptidase was selected as a no binding negative control for these experiments. Following the identification of the peptide core by NetMHCIIpan-4.0, the protein structures and identified peptide core were submitted to HPEPDOCK Server for peptide–protein molecular docking (Zhou et al., 2018). In brief, cleaned protein structures were used as receptors, and the peptide core sequence was used to generate 100 different conformers and a global sampling of binding orientations into the peptide binding domain of HLA-II receptors. Following docking, the peptide-HLA-II complexes with the highest complementarity were selected for receptor–peptide refinement in the HADDOCK Refinement Interface (Dominguez et al., 2003). Finally, the peptide-HLA complexes were analyzed for the formation of molecular interactions and binding energy using PLIP (Adasme et al., 2021) and PRODIGY (Honorato et al., 2021; Vangone and Bonvin, 2015).

### 6. Metagenomic analyses

#### Metagenomic sequencing

Metagenomic collection and sequencing has previously been detailed in (Zhernakova et al., 2016). In brief, participants collected and stored in the freezer their fecal samples directly at home. Fecal samples were collected on dry ice and transferred to the laboratory. Aliquots were stored at −80°C until further processing. The allPrep DNA/RNA Mini Kit (Qiagen; cat. 80204) was used for DNA isolation. DNA was sent to the Broad Institute (Cambridge, Massachusetts, USA) where library preparation and shotgun metagenomic sequencing were performed on Illumina HiSeq.

#### Metagenomic processing

Low-quality reads were discarded by the sequencing facility. Reads aligning to the human genome or to Illumina sequencing adapters were removed using default parameters using the KneadData pipeline (version 0.39). In short, this software uses Trimmomatic (Bolger et al., 2014) for adapter removal and quality trimming of reads and Bowtie2 (Langmead and Salzberg, 2012) for mapping and removal of reads mapped against the human genome (hg19). Taxonomy abundance estimation was then performed using MetaPhlan3 and default parameters (Beghini et al., 2021). Next, microbial relative abundance was transformed using additive log-ratios on the relative abundance table (adding ½ of minimal non-zero relative abundance to each cell in the table), with species geometric mean as denominator (center-log ratio). Bacteria not present in at least 10% of samples were discarded.

#### Microbiome-peptide association analysis

Co-occurrence between fecal microbiota and blood antibody–bound peptides was assessed using logistic regression analysis, while adjusting for the effects of age, sex and PhIP-Seq plate in 1,051 participants. In total, we analyzed the relation between 284 bacteria and 2,815 antibodies. Each antibody-bound peptide was modeled in generalized linear models as a response variable in a model including age, sex, PhIP-Seq plate and transformed bacterial abundance as predictors.

#### Microbiome meta-analysis

To increase the statistical power to detect associations between gut microbiota and blood antibodies, we combined the results of our cohort with the results derived from the 1000IBD cohort (n = 137, blood and fecal samples collected with <1 year difference) by performing a meta-analysis. We filtered out peptides not seen in at least 10 samples in both IBD and LLD cohorts. Heterogeneity coefficients (I^2^ and Cochran’s Q) were estimated per association. Meta-analysis was conducted by pooling summary statistics for both cohorts and under random and fixed-effects assumptions using the *meta* R package (v4.19-0) (Schwarzer and Others, 2007). FDR was estimated (Benjamini and Hochberg, 1995) from the resulting associations.

## Acknowledgments

We thank K. Mc Intyre for English editing.

The Lifelines Biobank initiative has been made possible by a subsidy from the Dutch Ministry of Health, Welfare and Sport; the Dutch Ministry of Economic Affairs; the University Medical Center Groningen; the University of Groningen and the Northern Provinces of the Netherlands. The authors wish to acknowledge the services of the Lifelines Cohort Study, the contributing research centers delivering data to Lifelines and all the study participants.

## Funding

The researchers participated in this project are supported by Netherlands Heart Foundation (IN-CONTROL CVON grants 2012-03 and 2018-27 to J.F. and A.Z.); the Netherlands Organization for Scientific Research (NWO) Gravitation Netherlands Organ-on-Chip Initiative to J.F. and C.W.; NWO Gravitation Exposome-NL (024.004.017) to J.F., A.K. and A.Z.; The Seerave foundation and the Netherlands Organization for Scientific Research to RKW; NWO-VIDI (864.13.013) and NWO-VICI (VI.C.202.022) to J.F.; NWO-VIDI (016.178.056) to A.Z.; NWO-VIDI (016.171.047) to I.J., NWO Spinoza Prize SPI 92-266 to C.W.; the European Research Council (ERC) (FP7/2007-2013/ERC Advanced Grant 2012-322698) to C.W.; ERC Starting Grant 715772 to A.Z.; ERC Consolidator Grant (grant agreement No. 101001678) to J.F.; and RuG Investment Agenda Grant Personalized Health to C.W.; A.R.B. [grant no. 17-57] and T.S. [grant no. 17-34] hold scholarships from the Junior Scientific Masterclass, University of Groningen. E.S. is supported by grants from the European Research Council, the Israel Science Foundation and by the Seerave foundation. T.V. gratefully acknowledges support from the Austrian Science Fund (FWF, Erwin Schrödinger fellowship J4256). I.J. and A.Z. were supported by a Rosalind Franklin Fellowship from the University of Groningen.

## Author contributions

Conceptualization: SA-S, ARB, SZ, JF, RKW, IJ, TV, SL

Methodology: SA-S, AK, SH, AJR, AVV, ARB, TS, SL, TV, SK, INK

Investigation: SA-S, ARB, AK, SH, AR, AVV, TS, TV, AZ, JF, IJ

Funding acquisition: SZ, JF, RKW, CW

Supervision: SZ, JF, RKW

Writing – original draft: SA-S, ARB

Writing – review & editing: All coauthors

## Declaration of interests

RKW acted as consultant for Takeda, received unrestricted research grants from Takeda, Johnson & Johnson, Tramedico and Ferring and received speaker fees from MSD, Abbvie and Janssen Pharmaceuticals. All other authors declare no competing interests.

## Supplementary Material

**Supplementary table 1:** Antibody-bound peptide general information. **1.1** Information from 2,815 analyzed peptides, including: database source, amino acid sequence, source protein name, source taxonomy, heritability estimate (H^2^), co-occurrence module belonging and consistency (after 4 years). **1.2** Left table. Summary of peptides belonging to each of the 22 modules with at least 10 peptides. Right table. General overview of the co-occurrence modules, their category, (1) same taxonomy, (2) ortholog protein, (3) unrelated taxonomy and structure, and the correlation of their *eigengenes* (P_Bonferroni_<0.05) **1.3** LLD phenotypes, exploratory statistics.

**Supplementary table 2.** Association analyses summary statistics. **2.1** Antibody-bound peptide among sample dissimilarity (Jaccard) analysis of variability, summary statistics (PERMANOVA, 2,000 permutations). **2.2** GWAS meta-analysis summary statistics (P < 5×10^−8^). **2.3** HLA associations meta-analysis summary statistics (P_Bonferroni_ < 0.05). **2.4** Microbiome taxonomic abundance associations summary statistics (P < 1×10^−3^). **2.5** Phenotype associations summary statistics (FDR < 0.05).

---

**Supplementary Figure 1.**
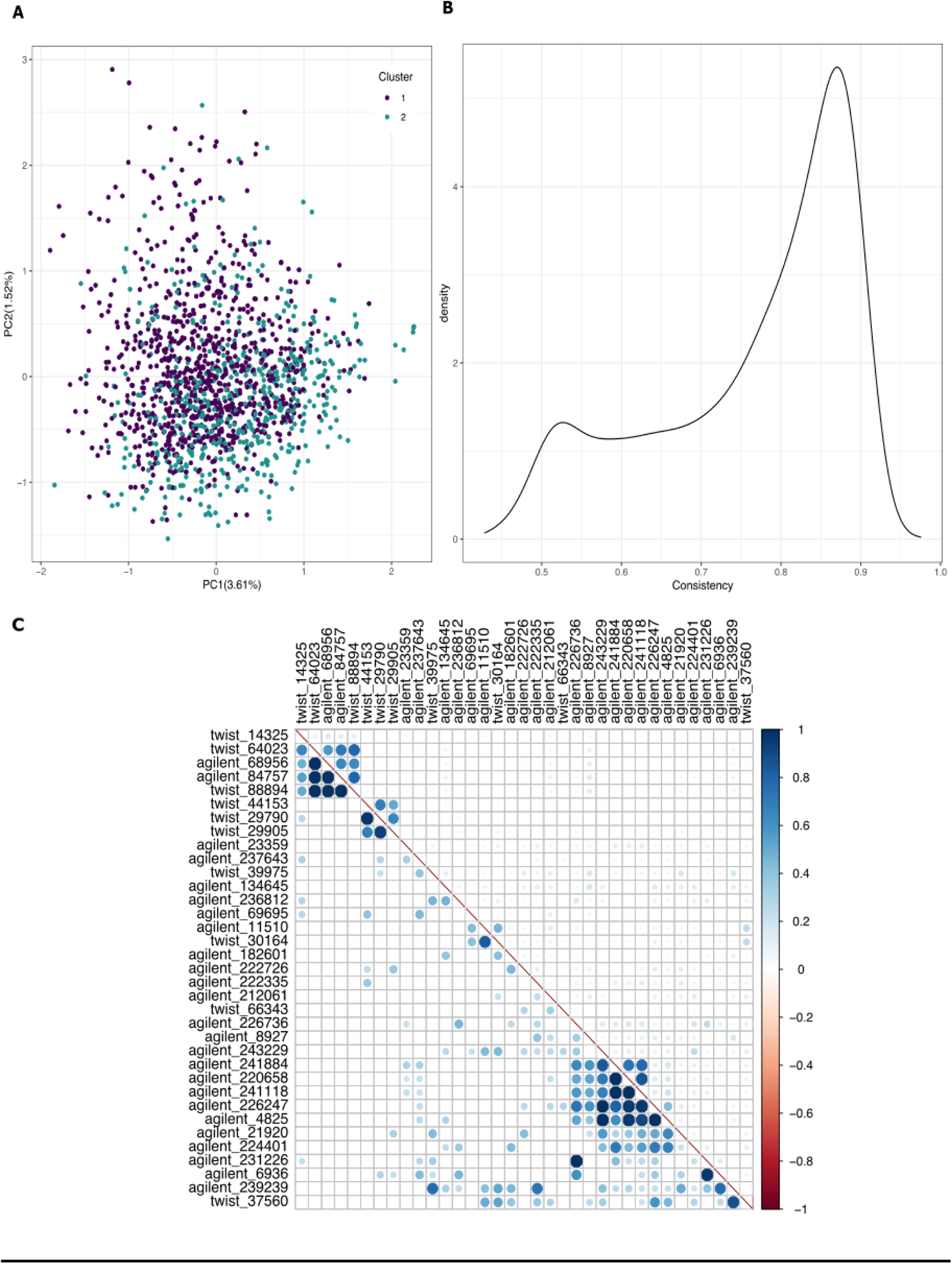
**A.** Antibody-bound peptide PCA after removal of 90 peptides belonging to CMV. **B.** Density of 2,815 antibody-bound peptide time consistency (same presence status in baseline as in follow-up) in 322 participants after 4 years. **C.** Correlation plot of highly heritable peptides (H^2^ ≥ 0.5). Lower triangle shows genetic correlation coefficient estimates. Upper triangle shows presence/absence Pearson’s correlation coefficients. Dot size and color indicate the strength of the correlation.

**Supplementary Figure 2.**
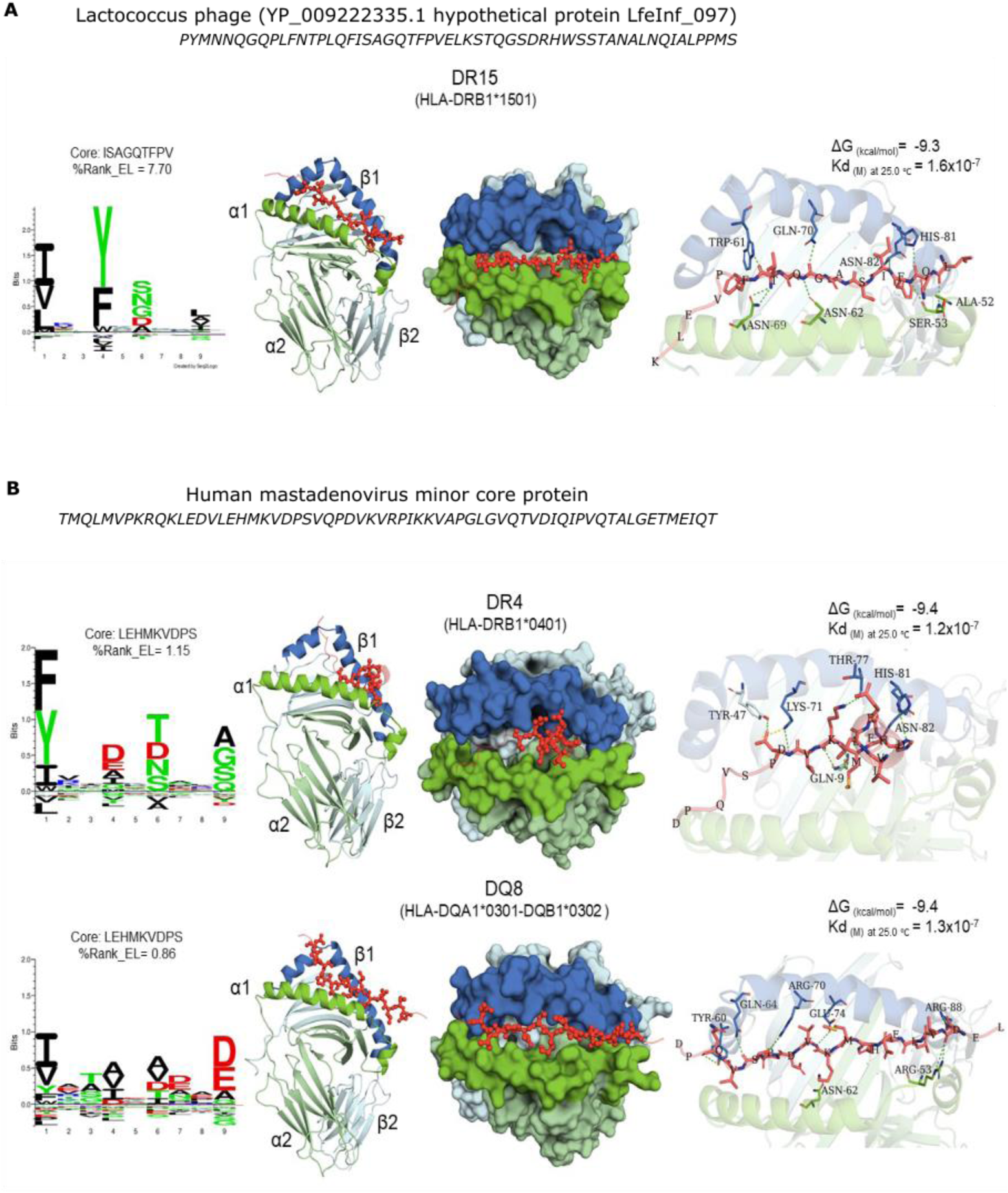
Peptide motif deconvolution map of **A.** DR15 and **B.** DQ8 and DR4 (amino acids code: negatively charged: red; positively charged: blue, polar uncharged: green, and hydrophobic: black) compared with the **A.** Lactococcus phage (YP_009222335.1 hypothetical protein LfeInf_097) and **B.** H*uman mastadenovirus* minor core protein. Peptide cores and percentage of elution score (%Rank_EL: strong binding ≤ 2.0, weak binding 2.0–10.0, no binding > 10) predicted by NetMHCIIpan-4.0 (Reynisson et al., 2020a) are shown. Predicted binding mode, polar molecular interactions (dashes, hydrogen bonds: green, salt bridges: yellow), binding energy and dissociation constant (Kd) of the Streptococcus agalactiae C5a peptidase peptide core (red cartoon and sticks) into HLA-II receptors (chain A in green and chain B in blue).

