## Supplementary figures and images for "Genetic, environmental and intrinsic determinants of the human antibody epitope repertoire"

### Supplementary Figure 1

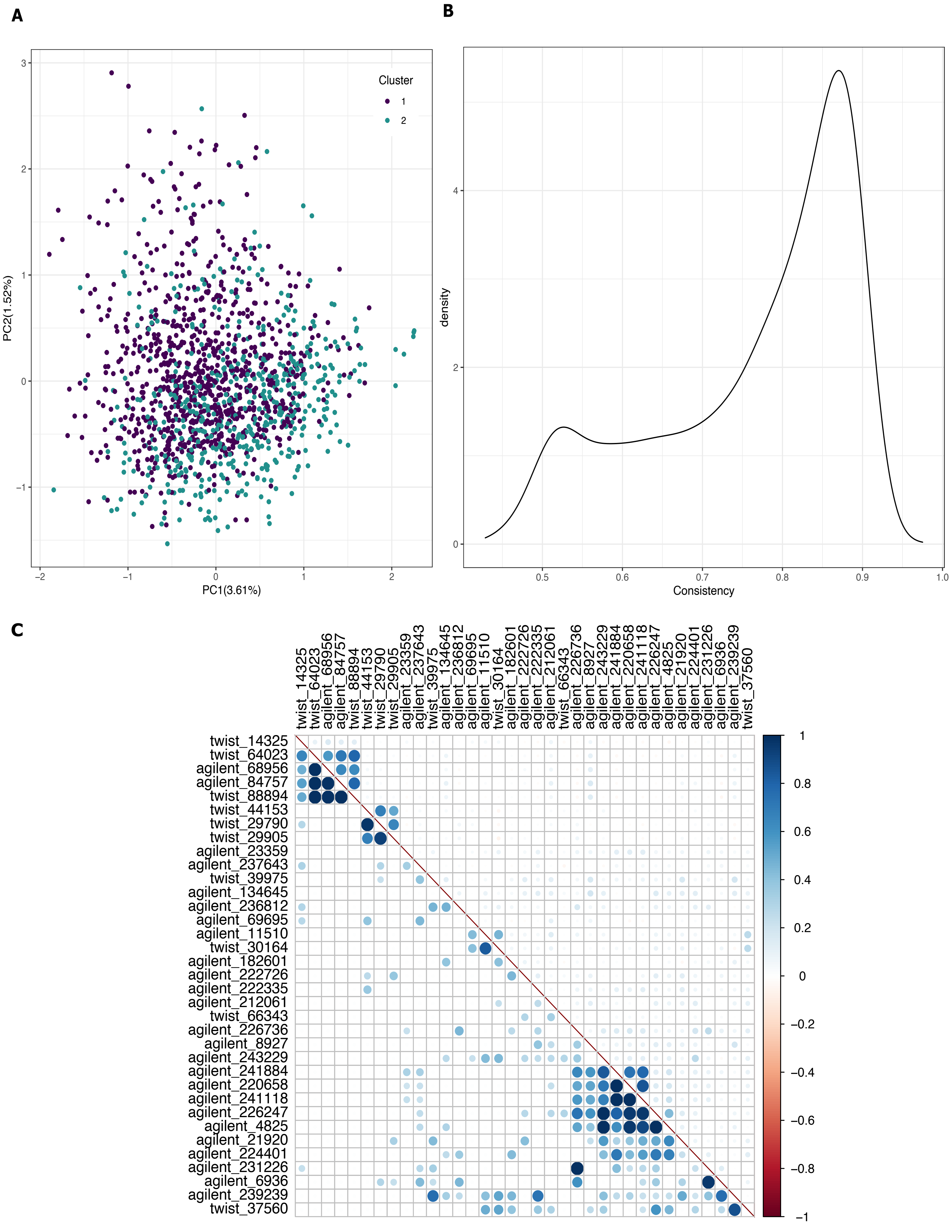

### Supplementary Figure 2

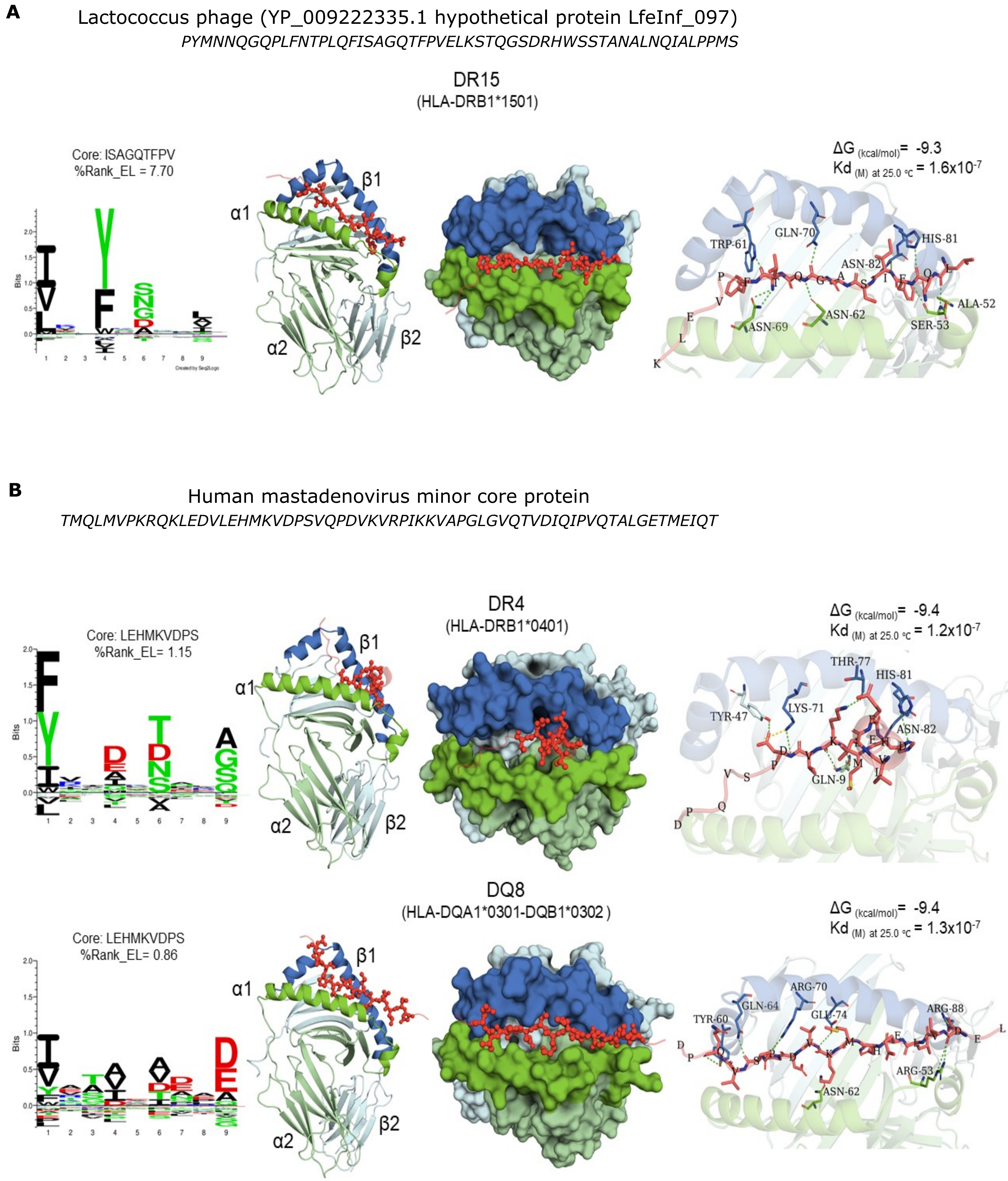
